## Supplementary figures for "Food intake enhances hippocampal sharp wave-ripples"

Supplementary Materials

Supplementary Figure 1.

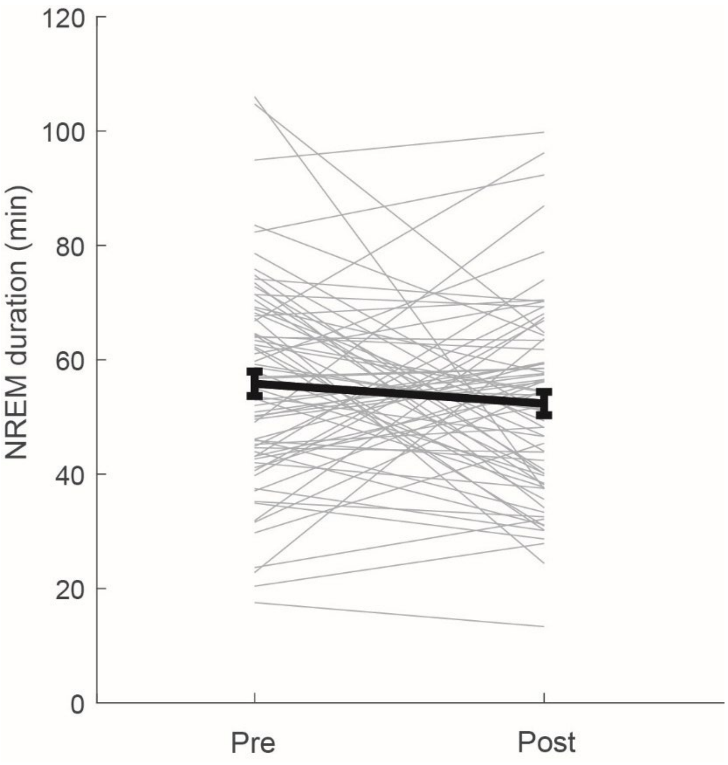


Comparison of NREM sleep durations in the Pre and Post sessions. Included are all session pairs in which animals were food restricted and received 0-1.5 g chow between the Pre and Post sleep sessions. Depicted is the total amount of time spent in NREM sleep within the 2 hour sleep session (wake and REM sleep excluded, all NREM bouts concatenated). NREM sleep amounts did not significantly differ in the Pre and Post sessions (P=0.1759, paired t-test).

Supplementary Figure 2.

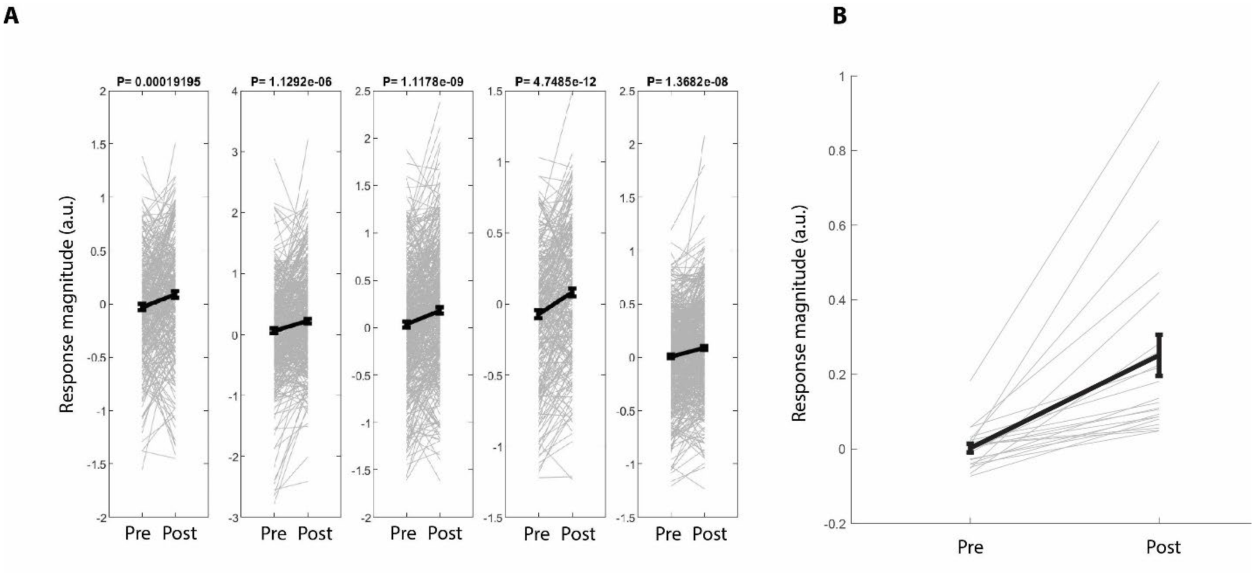


Additional quantification of SWR-triggered increase in LH GABAergic activity. A. Integrated fluorescence immediately before (“Pre”) and after (“Post”) SWRs. Each gray line corresponds to fluorescence magnitudes before and after a single SWR. Black line and bars indicate the pre/post average (means and standard error of the mean). Panels correspond to single sleep session in different animals. The p-value in the title of each plot corresponds to the significance of the SWR-triggered increase (corresponding to the values in main figure 5I). B. The SWR-averaged activity before and after SWRs from all sessions (each gray line represents the SWR-averaged values from one session). There was a significant SWR-triggered increase in LH GABA activity across sessions (P= 5.3337e-05, paired t-test).
